# Determining the shelf life of an erythropoietin alfa biosimilar GBPD002 through stability study

**DOI:** 10.1101/2023.08.05.552105

**Authors:** Kakon Nag, Mohammad Mohiuddin, Samir Kumar, Md. Maksudur Rahman Khan, Md. Enamul Haq Sarker, Bipul Kumar Biswas, Rony Roy, Md. Tarek Molla, Ratan Roy, Md. Emrul Hasan Bappi, Arifur Rahman, Sheik Rejaul Haq, Md. Shofiquzzaman Sarker, Priyanka Mollik Popy, Raisa Ferdaushi Mumu, Uttam Barman, Md. Shamsul Kaunain Oli, Md. Sadek Hosen Khoka, Sourav Sarker, Md. Firoz Alam, Naznin Sultana

**Affiliations:** Globe Biotech Limited, 3/Ka (New) Tejgaon I/A, Dhaka 1208, Bangladesh; R&D Management Solution Inc., Hamilton, Ontario L9C2V8, Canada

**Keywords:** Erythropoietin, stability study, shelf life, real-time stability, accelerated stability, GBPD002

## Abstract

GBPD002 is a recombinant human erythropoietin (rhEPO) produced by recombinant DNA technology using mammalian cell expression system. In this study, samples were analyzed according to current Good Laboratory Practice (cGLP) and regulatory guidelines to evaluate the quality of the product under the influence of variety of environmental factors in a time-dependent manner. Accelerated (25 ± 2 °C and relative humidity: 60 ± 5 %) and real-time (5 ± 3 °C) stability study were conducted up to 6 months and 12 months, respectively; samples were analyzed in every 3 months. After 12 months, real-time stability studies were performed with 6 months interval up to18 months, which to be continued up to 24 months. Appearance and pH were assessed using standard methods, and molecular weight was determined by Western blotting. Chromatographic identification and quantitative assays were performed by reversed-phase chromatography (RPC). High-molecular weight aggregatesand degradants were determined using size exclusion chromatography (SEC) and particle size distribution (PSD) analysis. Biofunctionality of the samples were evaluated by *in vitro* and *in vivo* bioassays. Bacterial endotoxin and sterility test were performed as safety parameters. All samples met the acceptance criteria, and the data were extrapolated using the regulatory guideline to determine the shelf life. The data revealed that the GBPD002 is stable up to 24 months.

## 1. Introduction

Erythropoietin (EPO) is a natural glycoprotein hormone. It is synthesized in kidney, and stimulates erythropoiesis by acting on erythrocyte precursors in bone marrow [1]. Human EPO (HuEPO) has an apparent molecular weight of 30.4 kDa, which consists of 165 amino acids and contains two disulfide linkages. The carbohydrate moiety of EPO contributes approximately 40% of its molecular weight and consists of 4 glycosylation sites, *viz.*, 3 N-glycans and a single O-glycan [2, 3]. Recombinant Human Erythropoietin (rHuEPO) is used for the treatment of anemic patients with chronic renal failure in whom the endogenous production of EPO is impaired. It is used in patients with chemotherapy-induced anemia [4], reduces the need for blood transfusions in patients who undergo surgery, and can also be used for patients at risk for perioperative transfusions with anticipated significant blood loss [5]. The glycoproteins are considered to be critical attributes of the biological functions [6]. At higher temperature, EPO initially dimerizes as well as degrades followed by high molecular weight (HMW) aggregates and low molecular weight (LMW) degradants [7, 8]. HMW aggregates as well as LMW degradants of therapeutic proteins may compromise their safety and efficacy [9, 10]. The primary concern is that the aggregates of therapeutic proteins may induce immune responses, which may have consequences ranging from reduction of product efficacy to patient fatality [11]. Therefore, the stability test is essential for determining and ensuring product quality, efficacy and safety by using various orthologous analytical techniques.

The stability study of a pharmaceutical product is a complex process that requires considerable time, expense, and scientific expertise to ensure formulation consistency and efficacy [12]. The pharmaceutical stability test is defined according to the International Conference of Harmonization (ICH) guidelines as a systemic test conducted in pharmaceutical products to demonstrate that drug quality is subjected to various environmental factors such as temperature, humidity and light for setting a drug test period or a pharmaceuticals shelf life and to recommend a good storage condition [13]. The USP defines the stability of the pharmaceutical product as ‘extension within certain limits’ and uses the same characteristics and attributes as it had when the products were made [14]. The identification, strength, purity, stability and pharmaceutical analysis of manufactured products are essential to assess shelf life or date of expiration to support the label claims [12,15]. Studies on stability must be carefully performed in compliance with guidelines regulated by ICH, World Health Organization (WHO), and similar bodies [16].

Stability tests are a routine operation used in the various phases of product development for drug substance and drug product. Early stages use accelerated stability tests to measure the type of aggregates as well as degradants those may be found during the long-term storage. The stability test ensures that the products remain in the market are fit for consumption throughout the declared shelf life [17]. In addition, it provides evidence about how the quality of a drug varies with time under the influence of a variety of environmental factors *e.g*., temperature, relative humidity (RH), light, pH and so on [12]. The higher complexities of high molecular weight proteins necessitate special requirements compared to low molecular weight drugs concerning analytical testing and quality assurance [18]. The efficacy of protein products depends not only on their amino acid sequence (primary structure) but also on their structures, which are determined by S-S linkages as well as hydrophobic and ionic interactions. A change in structure and conformation may significantly affect the potency of a protein drug or even lead to a complete loss of activity [19]. Therefore, stability analysis is not only important during the development and evaluation of a new biotechnological drug but is also a decisive aspect and regulatory parameter in the quality control of therapeutic proteins.

Recently, Globe Biotech Limited of Bangladesh has developed and manufactured an erythropoietin alfa biosimilar GBPD002, which was thoroughly characterized, went through successful clinical trial and got approval from the local regulatory authority for human administration [20, 21, 22, 23]. In this study, we investigated the shelf life of the product through accelerated and real-time stability study following regulatory framework. Relevant experiments and results are presented in this article.

## 2. Materials and Methods

### 2.1 Stability matrix design

A stability matrix is shown in Supplementary table 1, where all stability indicating parameters *e.g*., identification, pH, assay, *in vitro* and *in vivo* bio-assay, product and process-related impurities, appearance, sterility and endotoxin were included in the design in accordance with ICH Q1A, Q5C and Q6B guidelines [24].

### 2.2 Sampling

EPO concentrated solution was stock in sterile and endotoxin-free 2 D bag (Sartorius Stedim, Germany) and all batches of EPO were pre-filled in SyriQ BioPure^®^ 1mL syringe (Schott, Switzerland). For stability evaluation of GBPD002, Erythropoietin alfa concentrated solution or API-batches (Batch 01, Batch 02 and Batch 03) and GBPD002 finished product-batches (GBFP21013, GBFP21014, GBFP21015, GBFP21016, GBFP21017, GBFP21018, GBFP21019, GBFP21020, GBFP21021, GBFP21022, GBFP21023 and GBFP22024) were stored at accelerated storage condition (25 °C± 2 °C; 60% RH ± 5% RH) for 6 months and real-time storage condition (5 °C± 3 °C) for 24 months. Among these validation batches, there were 4 different strengths, which were 4000 IU/mL (GBFP21013, GBFP21014 and GBFP21015), 10000 IU/mL (GBFP21016, GBFP21017, GBFP21018,), 2000 IU/0.5 mL (GBFP21019, GBFP21020 and GBFP21021) and 5000 IU/0.5mL (GBFP21022, GBFP21023 and GBFP22024). A sampling plan is shown in Supplementary table 2, where samples were taken from both real-time and accelerated stability chambers for study [24].

#### Western blot

The samples were analyzed by Western blotting at reducing conditions using Eprex^®^ (Janssen Cilag, UK) as reference product. Analysis was done for identification test for molecular mass and immunochemical properties. All protein samples were loaded in Bolt 4 – 12% bis-tris plus gel (Thermo Fisher Scientific, USA) for separation by molecular size. Separated proteins were transferred onto iBot 2 PVDF membrane (Thermo Fisher Scientific, USA). Anti-Epo polyclonal antibody (Thermo Fisher, USA) was used as a primary antibody and Goat anti-rabbit (H+L) IgG HRP conjugated (Thermo Fisher, USA) used as a secondary antibody for capturing the target proteins. ECL chemiluminescence substrate (Thermo Fisher, USA) was used for reaction development and then captured the image by chemiluminescence colorimetric capture technique. Molecular size was determined by using novex sharp pre-stained protein marker (Thermo Fisher, USA). Novex^®^ ECL Chemiluminescent Substrate (Thermo Fisher Scientific, USA) for HRP was used to detect the signal and the imaging was done using Amersham Imager 600 RGB (GE Healthcare, USA).

#### RPC

The samples were analyzed for quantitative assay and chromatographic identification against Eprex^®^ using a reverse phase high performance liquid chromatography (RPC) method by Vanquish Core HPLC system (Thermo Fisher Scientific, USA) in a Hypersil GOLD C8 (Thermo Fisher Scientific, USA) column (2.1 × 100 mm, 175 Å pore size, 1.9 µm particle size.

Chromatographic control, data acquisition, and analysis were performed by using Chromeleon^TM^ 7 Chromatography Data System (CDS) software (Thermo Fisher Scientific, USA). The UV detection was operated at a wavelength of 280 nm. The mobile phase A (0.1% Formic acid in water) and mobile phase B (90% Acetonitrile + 0.1% Formic acid in water) were filtered using a 0.22-µm filter and degassed. The flow rate was 0.3 mL/min and the column were maintained at ambient temperature. The method run time was 25 minutes with a linear gradient of 100% (v/v) mobile phase B with mobile phase A. The injection volumes for both the test samples and references were 20 µL.

#### SEC

The samples were analyzed to determine impurities using size exclusion chromatography (SEC) method by an Ultimate 3000 UHPLC system (Thermo Fisher Scientific, USA) in an Acclaim SEC 300 (Thermo Fisher Scientific, USA) columns (4.6 × 300mm, 300 Å pore size, 5 µm particle size) using Eprex^®^ as reference product. The spectrophotometer UV detector was used at a wavelength of 280 nm. The mobile phase (1.5 mM potassium dihydrogen phosphate, 8.1 mM disodium hydrogen phosphate, and 0.4 M sodium chloride at pH 7.4) was filtered using a 0.22- µm filter and degassed. The flow rate was 0.3 mL/min and the column were maintained at ambient temperature. The method run time was 30 minutes. The injection volumes for both the test samples and references were 20 µL, Chromatographic control, data acquisition, and analysis were performed by using Chromeleon^TM^ 7 Chromatography Data System (CDS) software (Thermo Fisher Scientific, USA).

#### Particle size distribution (PSD)

Samples were prepared in 0.22 micron filtered 1× PBS (pH 7.2), and after stabilization at 20 °C for 20 min, analyzed in disposable plastic cuvette using a Zetasizer Nano ZSP (Malvern Panalytical Ltd., Malvern, UK) where respective buffers used as dispersant. The equipment was switched on minimum of 30–60 min before the experiment to stabilize the system. The refractive index (RI), viscosity, and dielectric constant of dispersion buffer (1× PBS, pH 7.2 at 20 °C) were considered 1.33, 0.88 cPs, and 79, respectively.

#### In vitro bioassay

The human erythroleukemia cell line TF-1 (ATCC, USA) were cultured in RPMI-1640 media (Thermo Fisher Scientific, USA) supplemented with 10% fetal bovine serum (FBS) (Thermo Fisher Scientific, USA), 2 mM L-glutamine (Thermo Fisher Scientific, USA), 1%(v/v) streptomycin (Thermo Fisher Scientific, USA). In addition, 12 ng/mL recombinant human granulocyte macrophage colony-stimulating factor (rhGM-CSF) (Thermo Fisher Scientific, USA) was included in the culture medium. 5×10^4^ cells/mL number of cells were seeded excluding FBS into 24 well plate with triplicate. Different similar concentration of erythropoietin preparation (0.4 IU/mL, 4 IU/mL, and 40 IU/mL) was used for both reference product and samples to demonstrate comparative study. Cells were maintained in 5% CO2 humidified incubator (Thermo Fisher Scientific, USA) at 37 °C and cells morphology were observed regularly. After 48 hours, cell proliferation was determined by the Trypan Blue exclusion method using EVOS^TM^ XL Core imaging system (Thermo Fisher Scientific, USA).

#### In vivo bioassay

In the first step of *in vivo* bioassay, 8–12 -week old 64 albino mice (weighing 17–21 g) were selected and randomly distributed into 8 cages. Randomization was done using the standard = RAND() function in Microsoft Excel (Microsoft, USA); treatments were given single-blinded in numerical order. There were eight mice in each cage. One cage administered with 0.05 mL buffer solution that was considered as placebo group other group treated with nothing considered as control group. Among the other six cages, three cages were distributed for sample-groups as treatment I (20 IU/mL), treatment II (40 IU/mL), treatment III (80 IU/mL) and other three cages were distributed for reference-groups as treatment I (20 IU/mL), treatment II (40 IU/mL), treatment III (80 IU/mL). Selected animals from each group received a pre-defined 0.05 mL subcutaneous injection at day 0, except control group. After the injections, on day 4, blood was collected from each mouse from the orbital sinus to determine the number of reticulocytes.

Whole blood was multiplied 500 times in the buffer used to make the colorant solution (thiazole orange) in order to prepare for analysis. The blood reticulocyte count was measured microfluorimetrically in a flow cytometer (BD FACSLyric^TM^, USA) after staining for 3–10 min. The percentage of reticulocytes is determined: number of cells/red fluorescence (620 nm).

Finally, potency was calculated by the usual statistical methods for a parallel line assay. All animal testing procedures were performed in compliance with the principles of laboratory animal care.

#### pH

pH was measured using calibrated SevenExcellence pH/ion multi-mode meter (Mettler Toledo, Germany). The sample volume was taken 1 mL for pH measurement.

#### Appearance

The contents of each syringe were examined for changes in color or clarity. This visual test was performed with the unaided eye under a laboratory light against a black and white background.

#### Sterility test

To screen for microbial (bacterial or fungal) contamination in stability samples, direct inoculation technique was used. Here, 20 µL of samples were inoculated in 3 mL of Tryptic Soya Broth (TSB) media for 14 days and then absorbance at 600 nm has been measured.

#### Endotoxin test

Bacterial endotoxin test of stability samples was performed using Pierce LAL Chromogenic Endotoxin Quantitation kit (Thermo Fisher Scientific, USA). All the steps were followed as per supplier’s instructions.

## 3. Results

### 3.1 Identification

Western blot was performed for molecular mass and immunochemical properties analysis. High intensity similar banding pattern were found at near around 34 kDa for both reference product and GBPD002stability samples (**Figure 1**).

**Figure 1:**
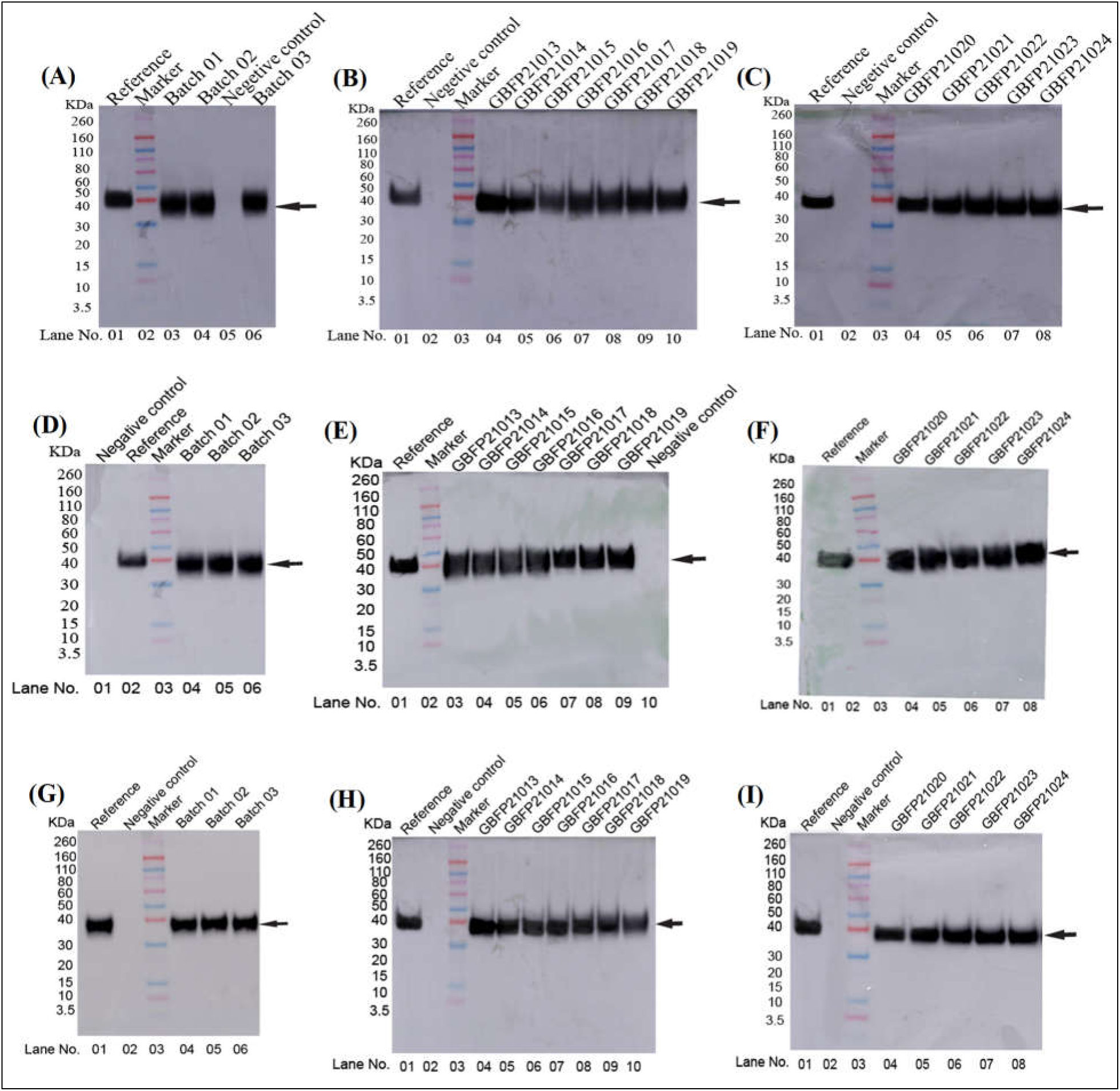
Analysis of molecular weight by Western blot. (A) real-time stability samples of API- batches. (B, C) real-time stability samples of finished product-batches. (D) accelerated stability samples of API-batches. (E, F) accelerated stability study samples of finished product-batches. (G) real-time stability samples of API-batches after 18 months. (H, I) real-time stability samples of finished product-batches after 18 months.

For the identification of the product, the retention time was found within specification limit (Retention time: 7.25 ± 0.20 minutes (**Figure 2**).

**Figure 2:**
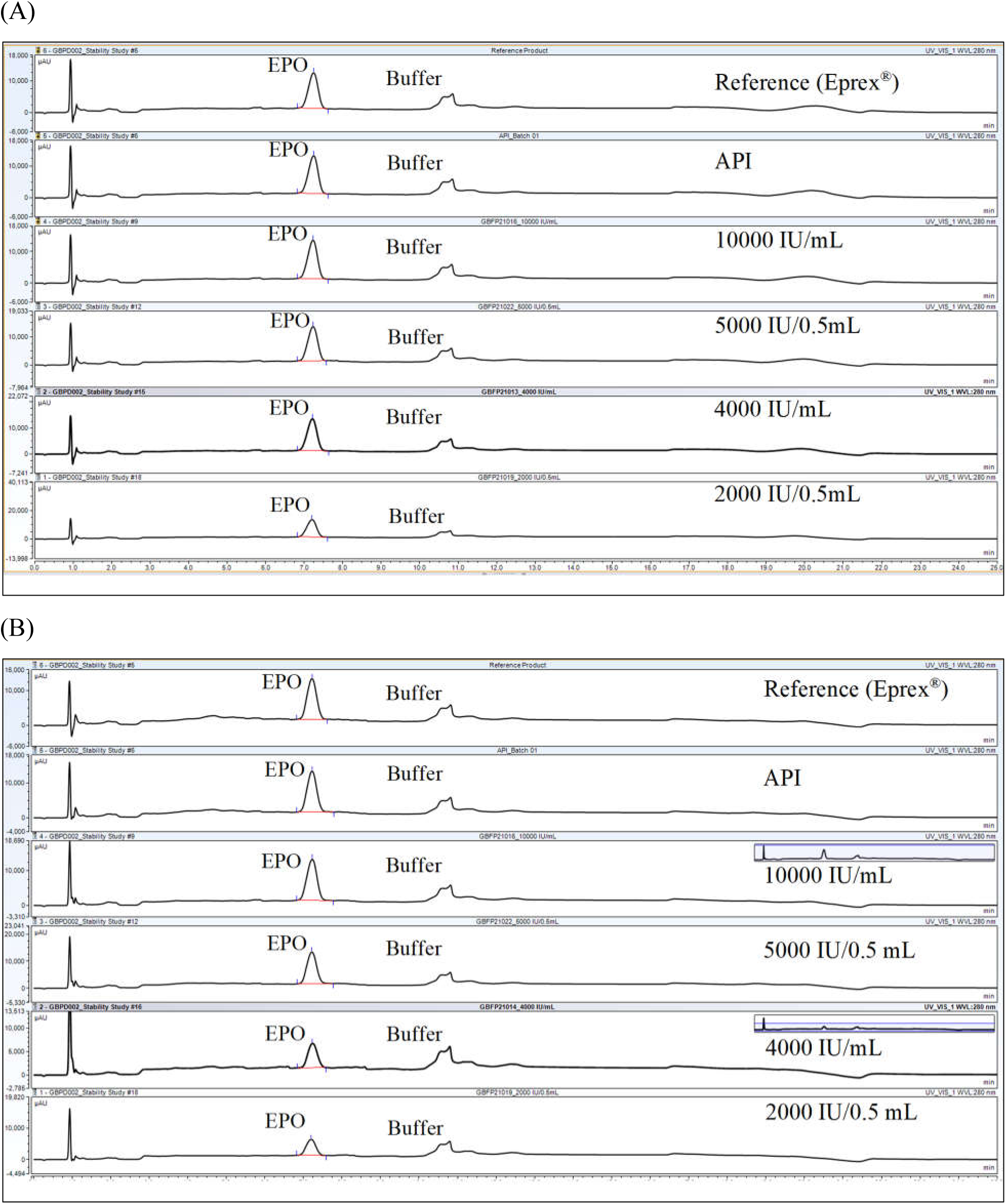
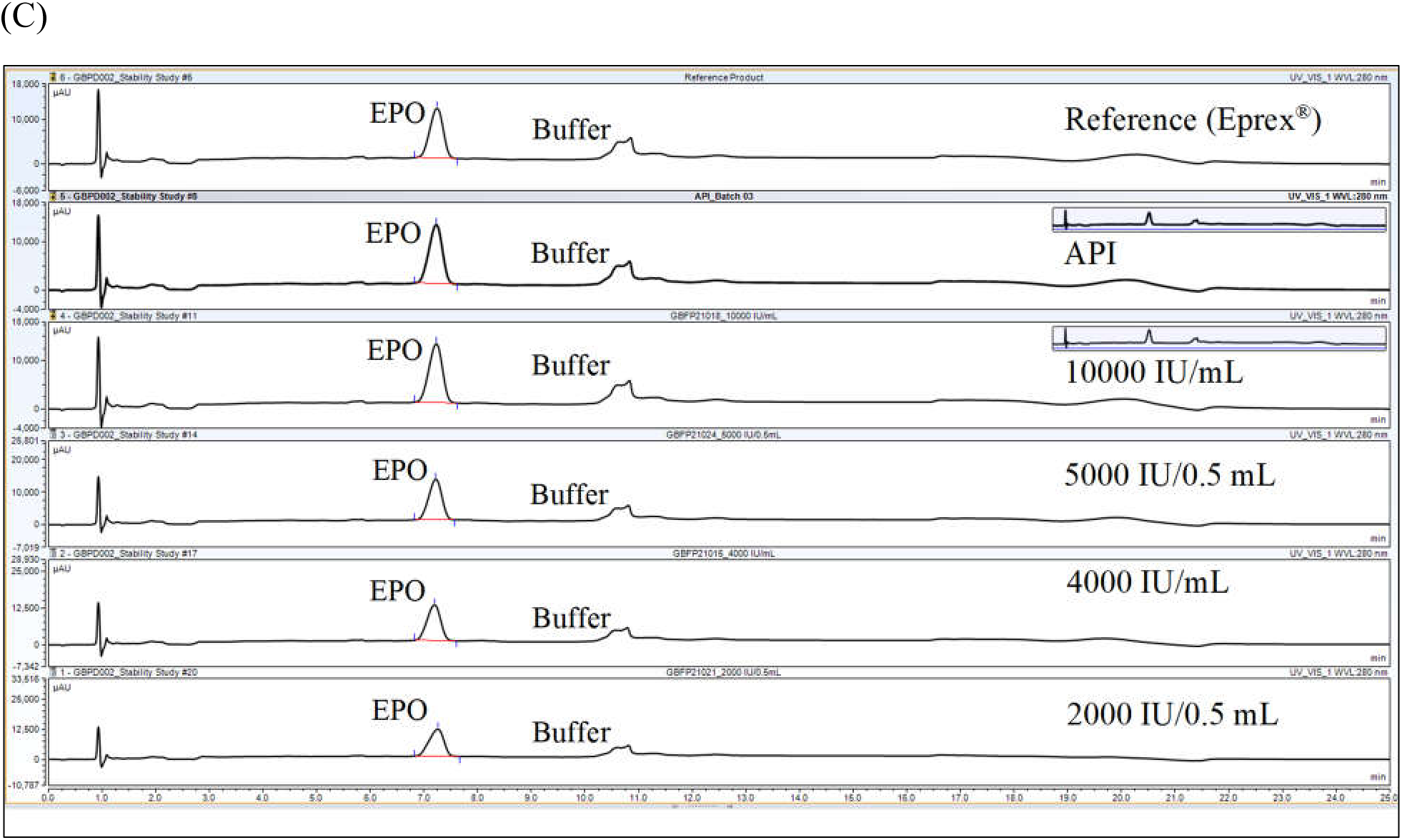
Identification of products for reference, API and finished products. (A) initial, (B) after 6 months at accelerated storage condition and (C) after 18 months at real-time storage conditions.

### 3.2 pH and Potency

The pH of the samples was found within specification (7.0 ± 0.3), no significant pH change was occurred among the all batches both for API (**Figure 3A and 3B**) and validation batches (**Figure 4A and 4B**). The potency of API (**Figure 3C and 3D**) and finished products (**Figure 4C and 4D**) were found stable within the acceptance limits (125 – 80%).

**Figure 3:**
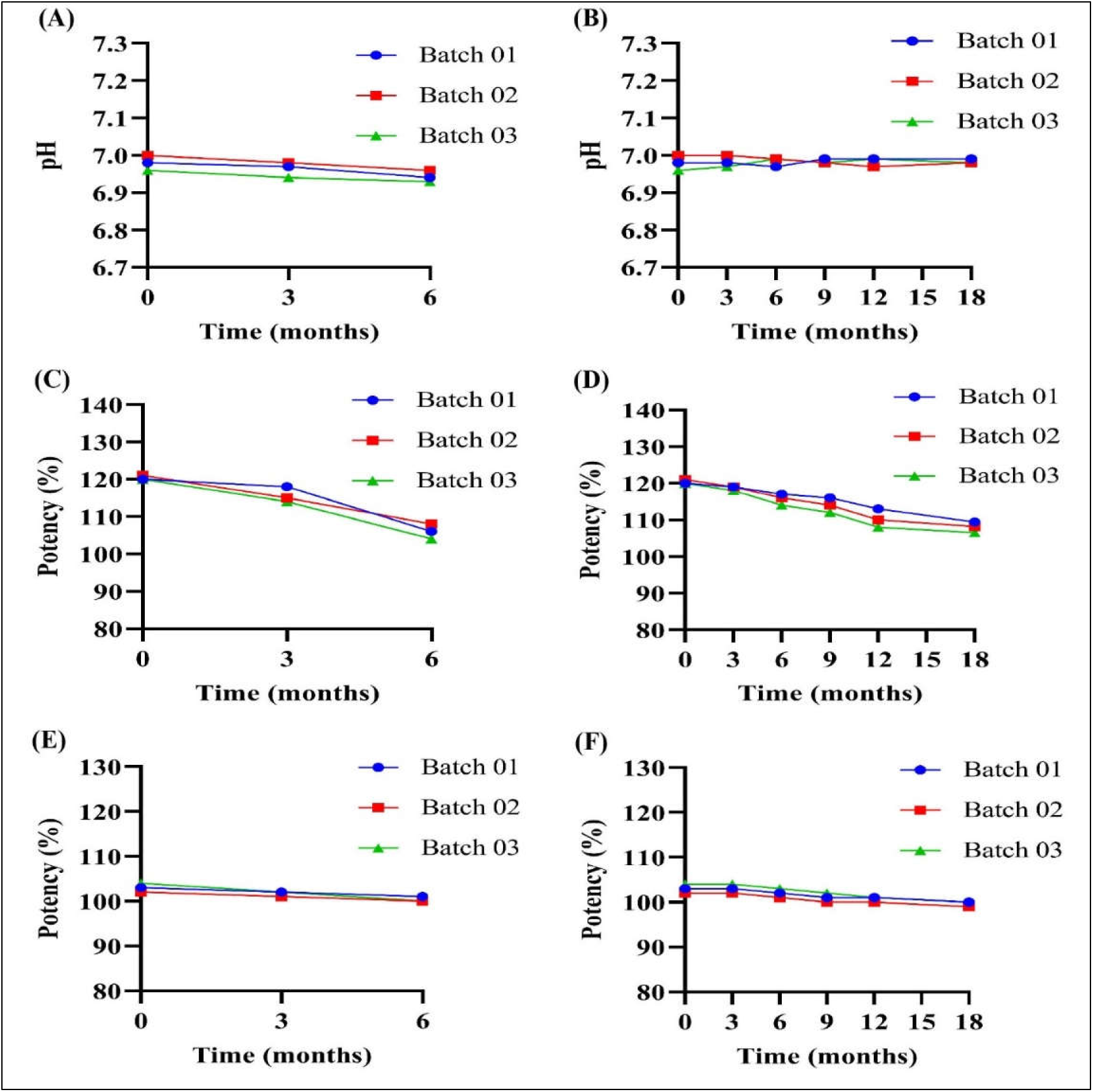
pH analysis (A, B), quantitative assay (C, D) and *in vivo* bioassay (E, F) of API- batches. (A, C, E) accelerated stability study, (B, D, F) real-time stability study.

**Figure 4:**
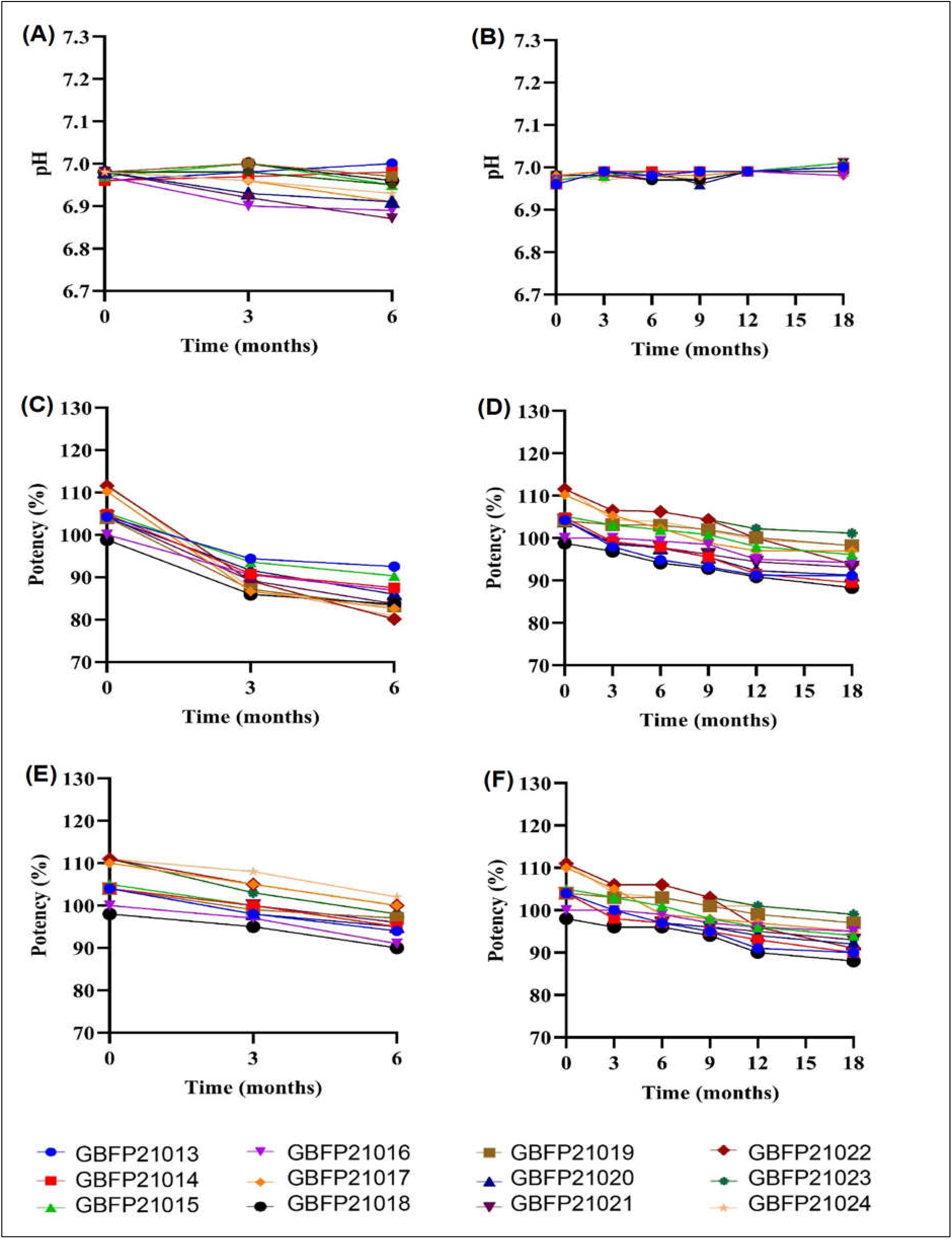
pH analysis (A, B), quantitative assay (C, D) and *in vivo* bioassay (E, F) of finished product-batches. (A, C, E) accelerated stability study, (B, D, F) real-time stability study.

### 3.3 Bio-assay

The *in vitro* dose-dependent cell proliferation was observed and found stable at different time- points of analysis for both API and finished products (**Figure 5 and 6**) and (**Supplementary table 3, 4, 5, 6 and 7**).

**Figure 5:**
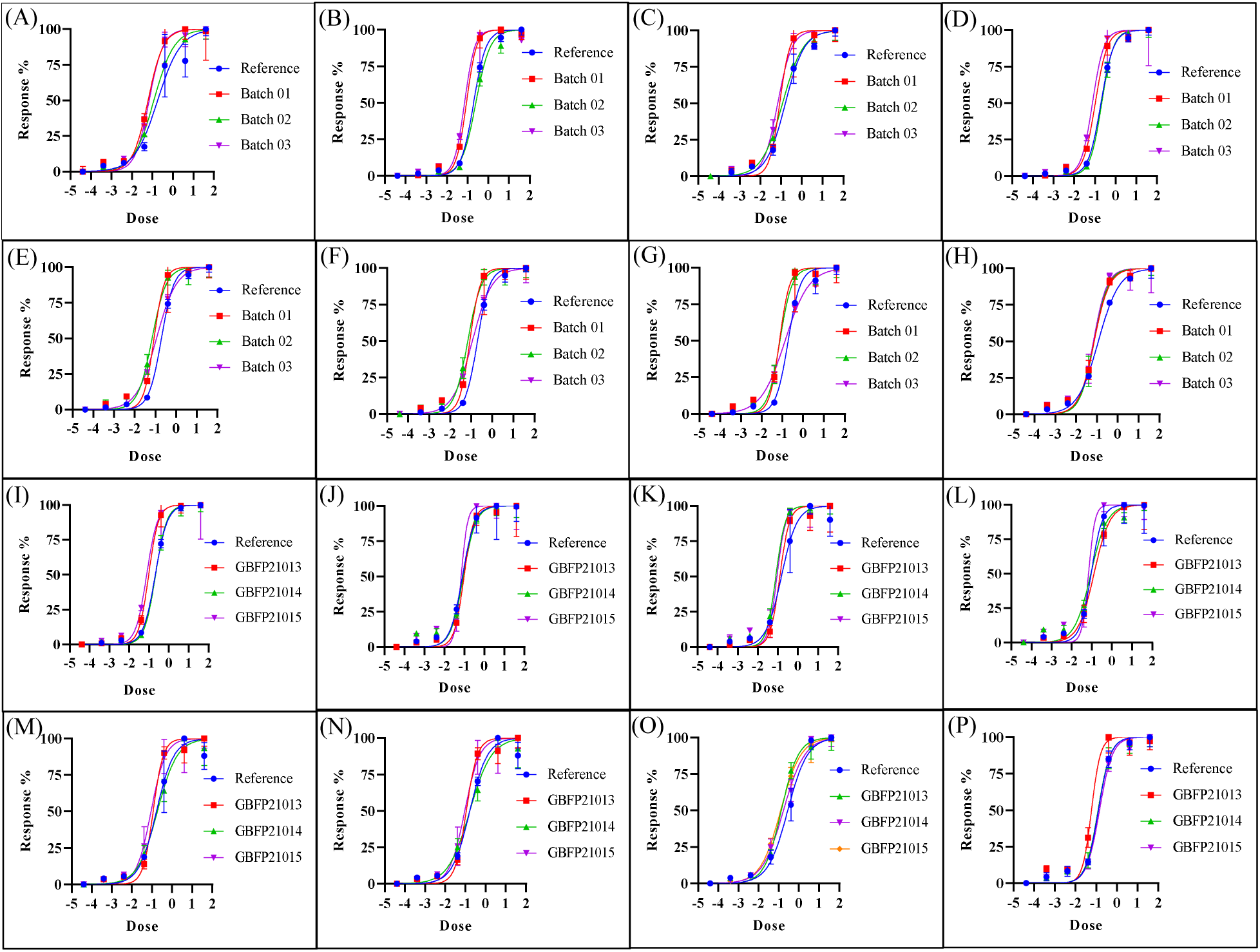
*In vitro* cell proliferation assay. For API GBPD002 (A) initial, (B, D) accelerated stability after 3 and 6 months respectively, (C, E, F, G and H) real-time stability study after 3, 6, 9, 12 and 18 months respectively. For finished product 4,000 IU/mL (I) Initial, (J, L): accelerated stability after 3 and 6 months respectively, (K, M, N, O, and P) real-time stability study after 3, 6, 9, 12 and 18 months respectively.

**Figure 6:**
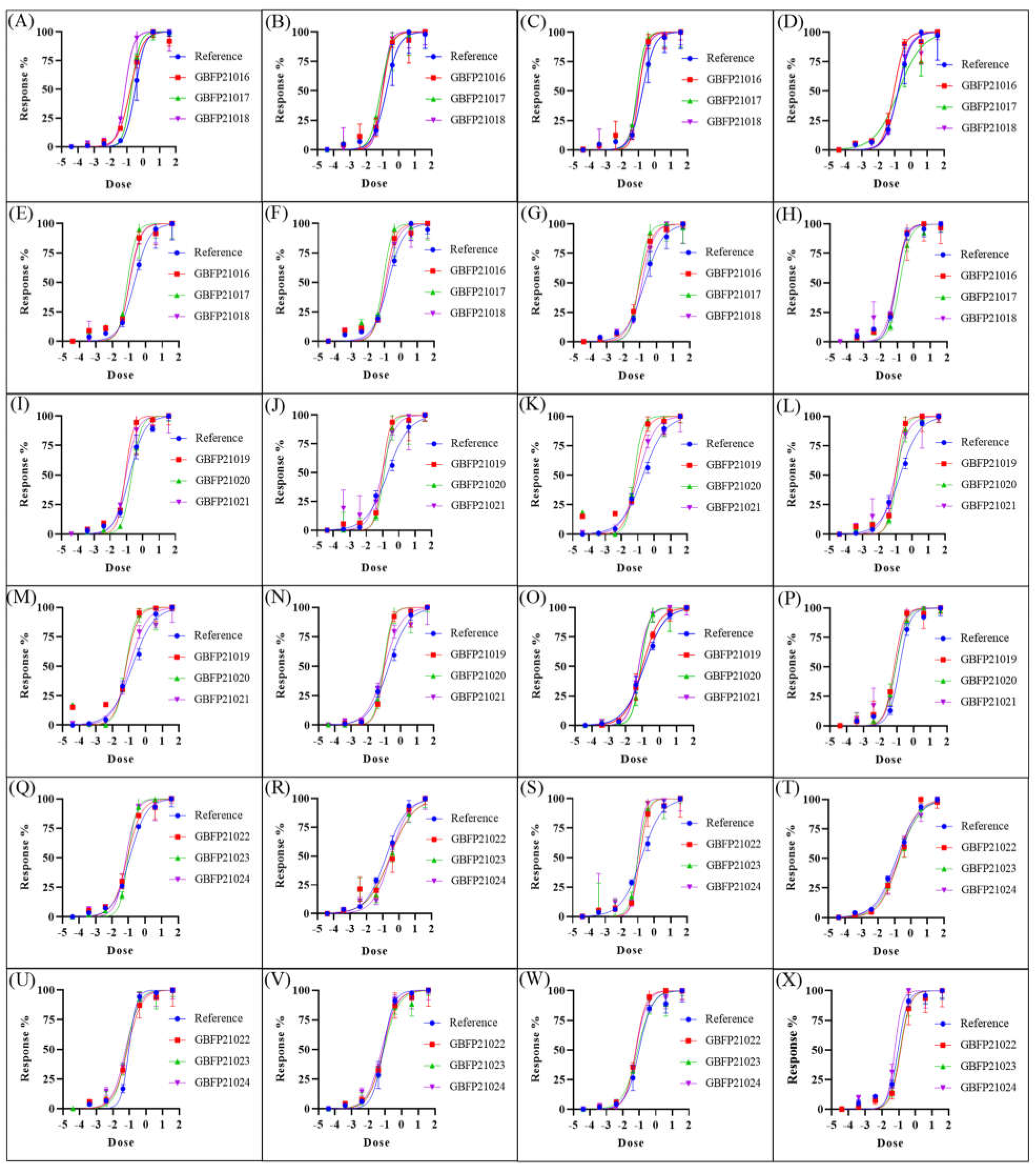
*In vitro* cell proliferation assay. For finished product 10,000 IU/mL (A) initial, (B, D) accelerated stability after 3 and 6 months respectively, (C, E, F, G and H) real-time stability study after 3, 6, 9, 12 and 18 months respectively. For finished product 2,000 IU/0.5 mL (I) Initial, (J, L): accelerated stability after 3 and 6 months respectively, (K, M, N, O, and P) real- time stability study after 3, 6, 9, 12 and 18 months respectively. For finished product 5,000 IU/0.5 mL (Q) Initial, (R, T): accelerated stability after 3 and 6 months respectively, (S, U, V, W, and X) real-time stability study after 3, 6, 9, 12 and 18 months respectively.

For *in vivo* bioassay, blood reticulocytes were counted for the biological assay (**Figure 7**).

**Figure 7:**
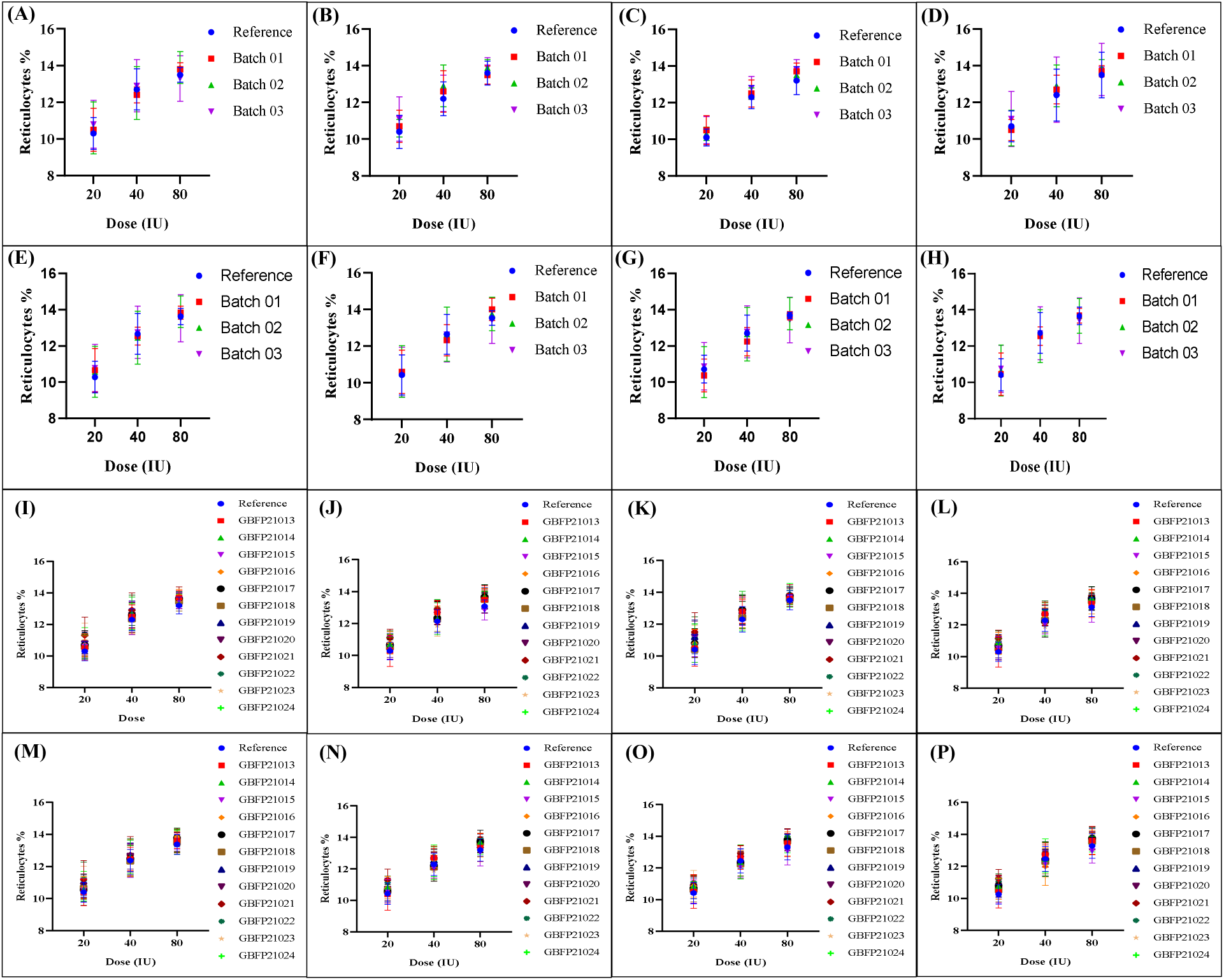
*In vivo* bioassay. For API (A) initial, (B, D) accelerated stability after 3 and 6 months respectively, (C, E, F, G and H) real-time stability study after 3, 6, 9, 12 and 18 months respectively. For finished products (I) Initial, (J, L): accelerated stability after 3 and 6 months respectively, (K, M, N, O, and P) real-time stability study after 3, 6, 9, 12 and 18 months respectively.

### 3.4 Product and process-related impurities

For product related impurities, aggregation was analyzed by SEC. A single product peak and blank peak was found for both API and finished products. No others aggregated and degraded peak was found (**Figure 8**).

**Figure 8:**
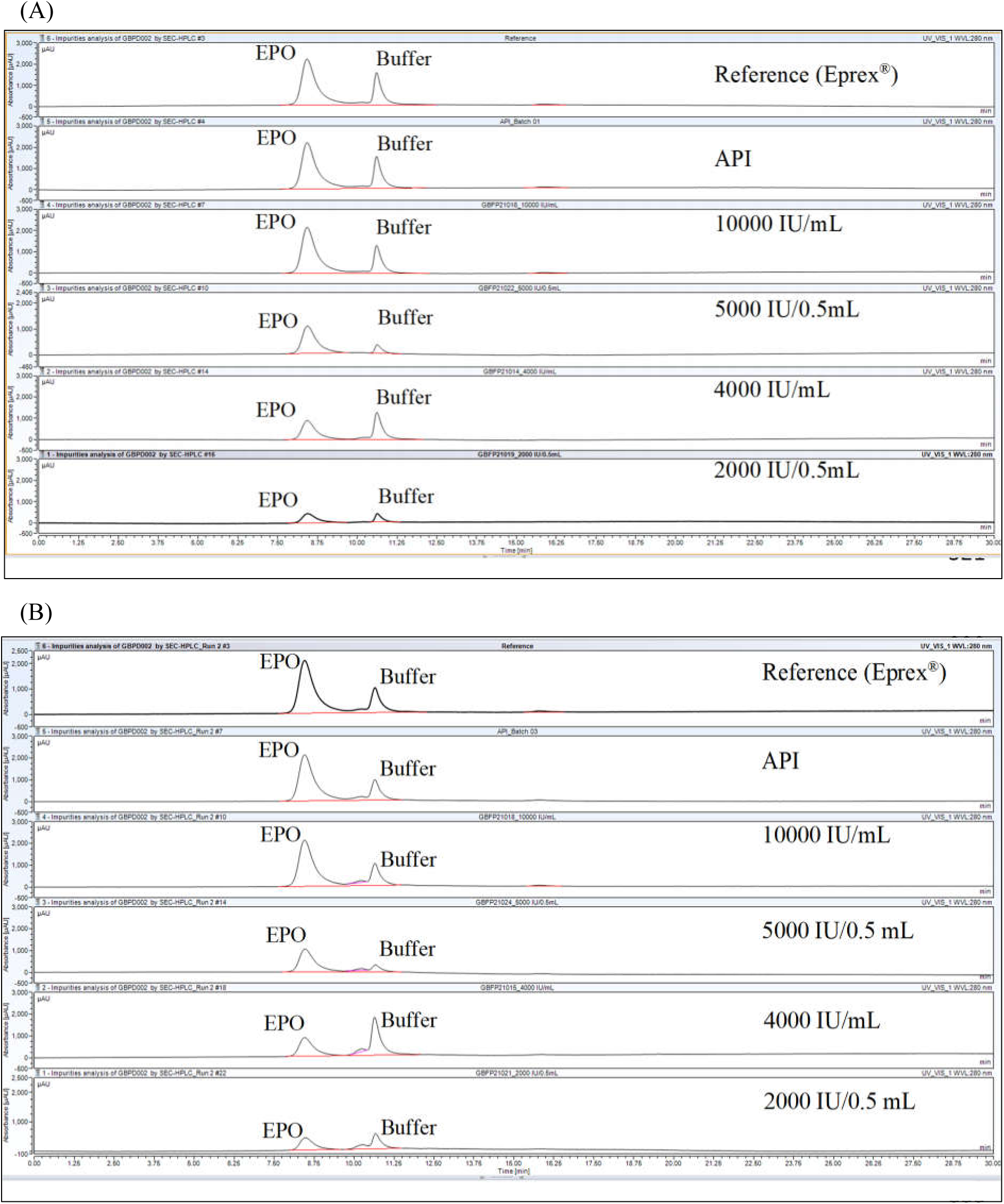
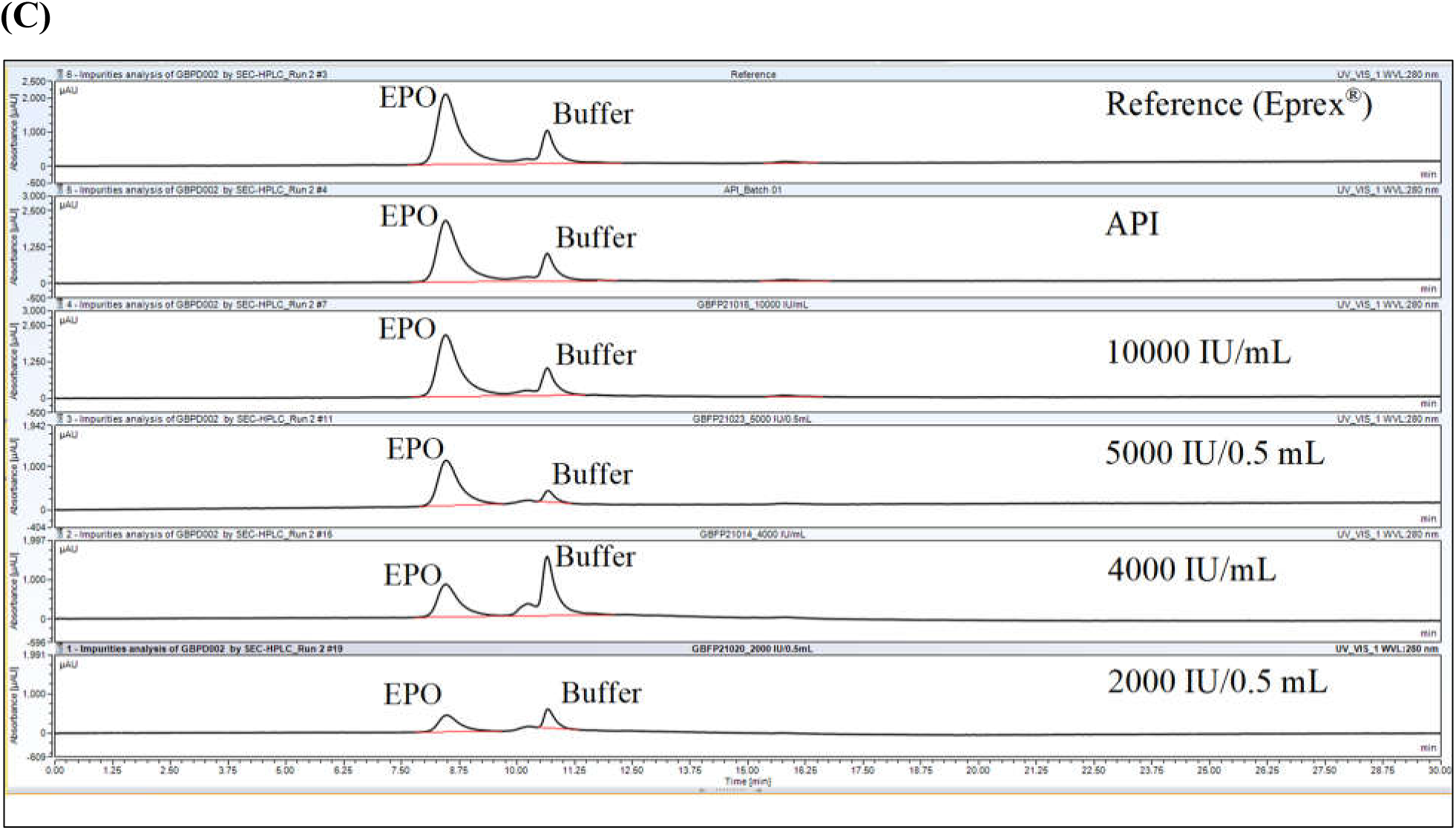
Aggregation and degradation analysis by SEC for reference, API and finished products. (A) representative data at initial, (B) representative data at 6 months for accelerated stability, and (C) representative data at 18 months for real-time stability.

In addition, aggregation and degradation was observed by PSD analysis where particle size of reference, API and finished products showed no aggregates and degradants was found (**Figure 9 and 10**).

**Figure 9:**
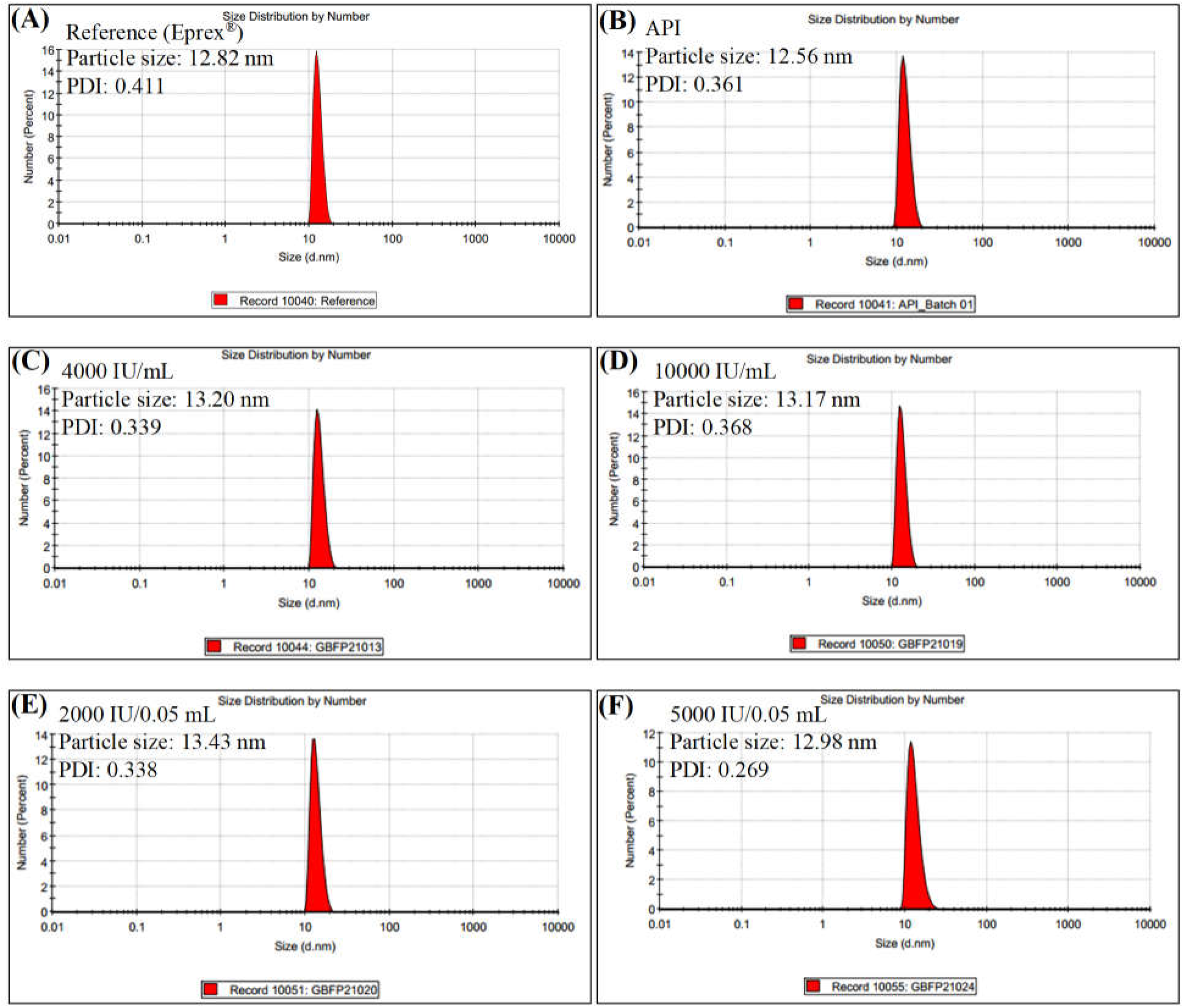
Particle size distribution analysis at initial point. Representative data for reference (A), API (B), and finished products (C, D, E and F).

**Figure 10:**
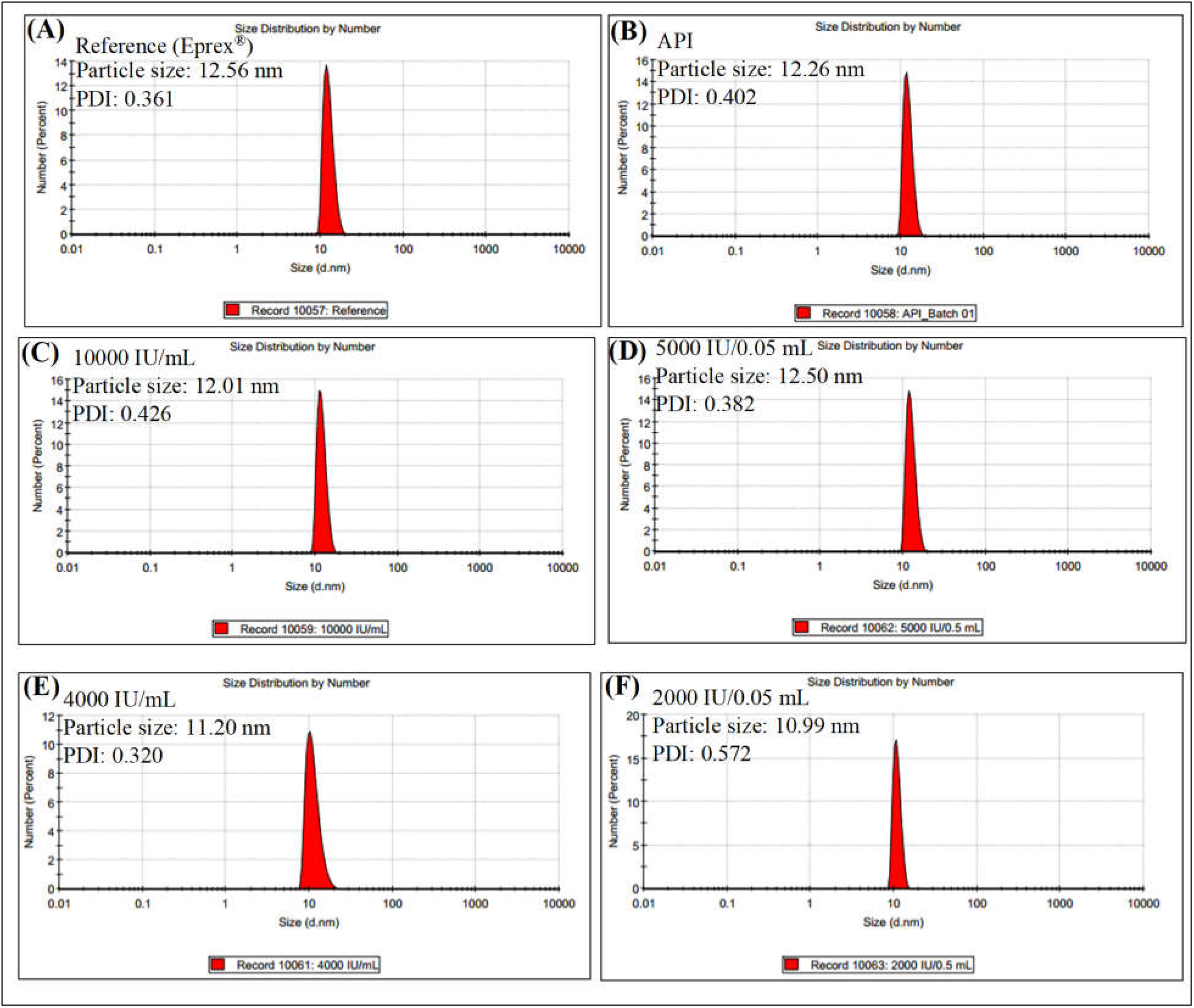
Particle size distribution analysis after 18 months at real-time storage conditions. Representative data for reference (A), API (B), and finished products (C, D, E and F).

### 3.5 Appearance, sterility and endotoxin test

During the stability study, there was no change in the appearance, color or clarity of all the batches sample. No growth was found for microbial sterility testing, and the endotoxin values were found below the specification limit and 2.5 EU/mL for each batch.

## 4. Discussion

The stability testing of biotechnological/biological products are highly recommended in ICH, WHO, and other regulatory guidelines because these productsaremostly sensitive tovarious environmental factors e.g., temperature, oxidation, light, ionic content etc. Hence, maintaining the biological activity as well as avoiding degradation and aggregation, strictconditions for storage are usually necessary and stability studies should be carried out in a scheduled manner [24]. In addition, the critical quality attributes (CQA) i.e., potency, purity and quality analysis of drug substance (bulk) and drug product (finished dose) in final container closure system are crucial to monitor the stability of a product in a pre-programmed matrix design [25].

The stability of a pharmaceutical productensures that the product in recommended storage and shelf-life conditions is safe and effective throughout its shelf life, provides evidence on how the quality of the drug substances or drug product changes over time under the influence ofenvironmental factors, and also establishes a retest or shelf life for drug substance or drug product and recommend the storage conditions [26].Therefore, the stability test should be conducted based on scientific principles and comprehension of current regulatory requirements [27, 28, 29]. Here, both real-time and accelerated studies were performed to obtain stability data.This stability testing of both API and finished products (GBPD002) included identification testing (Western blot: molecular size, and RPC: peak retention time), potency testing (RPC: assay, *in vitro* bio-assay: cell proliferation*, in vivo* bioassay: RET count), purity testing (SEC: aggregation and degradation, sterility: microbial contamination, LAL test: bacterial endotoxin) and other parameters (Appearance: visual observation, pH: ions). The appearance of all samples for both API and finished products of GBPD002 along with reference product were found within acceptance limits throughout the stability period. Approximately 34 kDa band in Western blotting confirmed that molecular mass of erythropoietin alfa in both API and finished products are stable up to 18 months at real-time storage and up to 6 months accelerated storage conditions. Up to 18 months, neither of the products show any degradation or aggregation in formulations at real-time storage. After 6-month, both API and finished products were found in degrading mode at accelerated storage conditions (**Figure 1D, 1E, and 1F**). As orthogonal methods for the identification of molecular mass, the retentions of erythropoietin alfa in RPC method were also investigated for both API and finished products. The retention times (7.25 ±0.20 minutes) of erythropoietin alfa in both formulations were found within specification limits in RPC (**Figure 2A** – **C**). The pH was also investigated, and all were found within acceptance limits (7.0 ± 0.3) in both at both real-time (5 ± 3 °C) and accelerated storage conditions (25 ± 2 °C) up to 6 months. After 6 months, both API and finished products were also found within specification limits up to 18 months. The trend line was shown similar for both API and finished products (**Figure 3A** – **B and Figure 4A** – **B**). We also investigated the assay quantity of erythropoietin alfa by RPC method at real time (from initial to 18 months) storage conditions and up to 6 months for accelerated storage conditions in both API and finished products. The trend line of assay quantity of erythropoietin alfa at accelerated (**Figure 3C and 4C**) and real time (**Figure 3D and 4D**) storage conditions was observed to check the similarity of assay quantity of all delivered doses found within specification limits (80–125%) in both API and finished products. The cell proliferation for both API and finished products was observed based on different doses *in vitro* at real time (from initial to 18 months) storage conditions and up to 6 months for accelerated storage conditions. The dose response curve of erythropoietin alfa at accelerated and real time storage conditions (**Figure 5 and 6)** was observed to check the similarity of cell proliferation of all delivered doses in both API and finished products. The effective concentration (EC50) was extrapolated from dose response curve and the EC50value were found similar for both API and finished products (**Supplementary table 3** – **7)**. The reticulocytes counts were observed based on 3 different doses of erythropoietin alfa *in vivo*at real time (from initial to 18 months) storage conditions and up to 6 months for accelerated storage conditions for both API and finished products. The reticulocytes count in mice after injection at accelerated and real time storage conditions (**Figure 7)** was observed to check the similarity of reticulocytes counts of all doses and found comparable for both API and finished products. In addition, we also investigated the potency of erythropoietin alfa *in vivo* at real time (from initial to 18 months) storage conditions and up to 6 months for accelerated storage conditions for both API and finished products. The trend line of potency of erythropoietin alfa at accelerated (**Figure 3E and 4E**) and real time (**Figure 3F and 4F**) storage conditions was observed to check the similarity of potency of all doses found within specification limits (80–125%), and found comparable for both API and finished products.

We also investigated the aggregation and degradation (product-related impurities) of erythropoietin alfa by validated SEC method at real time (18 months) storage conditions and accelerated storage conditions (6 months) for both API and finished products. No aggregates and degradants were found in both API and finished products. No additional peak other than the baseline reference of blank injection was observed for aggregates and degradants in both API and finished products and found similar (**Figure 8A** – **C**). As orthogonal analysis, particle size distribution was observed and found within acceptance limits (12 ± 5 nm) for initial and 18 months at real-time storage conditions (**Figure 9 and Figure 10**). The microbial contamination was investigated for API and finished products at real time and accelerated storage conditions.

No microbial growth was found in API and finished products. The bacterial endotoxin limit was found within specification limit (< 20 EU/mL) at real-time and accelerated storage conditions for both API and finished products.

## 5. Conclusion

The accelerated stability study revealed that the test product (GBPD002) is stable within the acceptance limits up to 6 months, and therefore, the shelf life of the product is claimed 24 months based on the data extrapolation principle. Real-time stability data revealed that the test product (GBPD002) is stable up to 18 months with respect to quality, safety and efficacy at 5 ± 3°C. The study ensures the conformity of quality product (GBPD002, brand name: **GB***poietin^®^*) for the stakeholders within the claimed shelf life. The next scheduled time-point for sample analysis at 24-month shall reveal the validity of the total time-span for the claimed shelf life.

## Author contributions

Conceptualization and study plan: Kakon Nag and Naznin Sultana; Test product manufacturing and evaluation: Samir Kumar, Md. Enamul Haq Sarker, Md. Bipul Kumar Biswas, Ratan Roy, Maksudur Rahman Khan, Rony Roy, Md. Tarek Molla, Arifur Rahman, Sheik Rejaul Haq, Md. Shofiquzzaman Sarker, Priyanka Mollik Popy, Raisa Ferdaushi Mumu, Uttam Barman, Md. Shamsul Kaunain Oli, Md. Sadek Hosen Khoka, Sourav Sarker, Md. Firoz Alam, Md. Emrul Hasan Bappi and Mohammad Mohiuddin; manuscript writing and editing: Kakon Nag, Naznin Sultana, Mohammad Mohiuddin, Samir Kumar, Md. Tarek Molla; All authors have read and agreed to the manuscript.

## Ethical statement

The experiments on TF-1 human cell line was approved by Internal Ethical Clearance Board (IECB), Globe Biotech Limited, Bangladesh (protocol number: GB/EC/18/001 and approval date: 15 February 2018). The study plan and procedures for *in vivo* experiments were approved by the internal ethical review board (IECB-PCS: Internal Ethical Clearance Board for Pre- Clinical Study) of Globe Biotech Limited, which is complied with the local and international regulation.

## Acknowledgements

The study was funded by Globe Biotech Limited. We thank Md. Harunur Rashid, the chairman of Globe Pharmaceuticals Group of Companies, Ahmed Hossain, Md. Mamunur Rashid, Md. Shahiduddin Alamgir and Abdullah Al Rashid, the directors of Globe Pharmaceuticals Group of Companies for their continuous support and encouragement. We also thank Dibakor Paul, Mithun Kumar Nag, Zahir Uddin Babor, GM Sajib Hassan, and Mijanur Rahman for their support for information and facility management system.

## Availability of data and materials

The data that support the findings of this study are available within the article and its Supplementary information file, or are available from the corresponding author upon reasonable request.

## Consent for publication

Not applicable.

## Supplementary file

Supplementary information.

## References

1. Jelkmann, W. Erythropoietin: Structure, Control of Production, and Function. Physiological Reviews 1992, 72 (2), 449–489. https://doi.org/10.1152/physrev.1992.72.2.449.

2. O’Connor, S. E.; Imperiali, B. A Molecular Basis for Glycosylation-Induced Conformational Switching. Chemistry & Biology 1998, 5 (8), 427–437. https://doi.org/10.1016/S1074-5521(98)90159-4.

3. Maruyama, K.; Miyata, K.; Yoshimura, A. Proliferation and Erythroid Differentiation through the Cytoplasmic Domain of the Erythropoietin Receptor. Journal of Biological Chemistry 1994, 269 (8), 5976–5980. https://doi.org/10.1016/S0021-9258(17)37558-0.

4. Egan, C.; Galli, L.; Ricci, C. Epoetin Beta for the Treatment of Chemotherapy-Induced Anemia: An Update. OTT 2015, 583. https://doi.org/10.2147/OTT.S77497.

5. McGirr, A.; Pavenski, K.; Sharma, B.; Cusimano, M. D. Blood Conservation in Neurosurgery: Erythropoietin and Autologous Donation. Can. J. Neurol. Sci. 2014, 41 (5), 583–589. https://doi.org/10.1017/cjn.2014.14.

6. Li, H.; d’Anjou, M. Pharmacological Significance of Glycosylation in Therapeutic Proteins. Current Opinion in Biotechnology 2009, 20 (6), 678–684. https://doi.org/10.1016/j.copbio.2009.10.009.

7. Arakawa, T.; Philo, J. S.; Kita, Y. Kinetic and Thermodynamic Analysis of Thermal Unfolding of Recombinant Erythropoietin. Bioscience, Biotechnology, and Biochemistry 2001, 65 (6), 1321–1327. https://doi.org/10.1271/bbb.65.1321.

8. Matejtschuk, P.; Duru, C.; Malik, K. P.; Bristow, A. F.; Costanzo, A.; Burns, C. J. Development of a Stable Chemically Cross-Linked Erythropoietin Dimer for Use in the Quality Control of Erythropoietin Therapeutic Products. Anal Bioanal Chem 2019, 411 (13), 2755–2758. https://doi.org/10.1007/s00216-019-01768-4.

9. Mitsudome, T.; Moribe, M.; Obayashi, Y.; Uchiyama, A.; Aono, M. Influence of Low-Molecular-Weight Aggregates on Aggregate Growth Kinetics and Physical Properties of Solid-State Proteins during Storage. European Journal of Pharmaceutics and Biopharmaceutics 2020, 146, 10–18. https://doi.org/10.1016/j.ejpb.2019.11.004.

10. Roberts, C. J. Therapeutic Protein Aggregation: Mechanisms, Design, and Control. Trends in Biotechnology 2014, 32 (7), 372–380. https://doi.org/10.1016/j.tibtech.2014.05.005.

11. Chang, Seong-Hun; Kim, Hyun-Jung; Kim, Chan-Wha. Analysis of the Structure and Stability of Erythropoietin by PH and Temperature Changes Using Various LC/MS. Bulletin of the Korean Chemical Society 2013, 34 (9), 2663–2670. https://doi.org/10.5012/BKCS.2013.34.9.2663.

12. F, Z.; R, R.; K, S. A REVIEW ON STABILITY TESTING GUIDELINES OF PHARMACEUTICAL PRODUCTS. Asian J Pharm Clin Res 2020, 3–9. https://doi.org/10.22159/ajpcr.2020.v13i10.38848.

13. Kommanaboyina, B.; Rhodes, C. T. Trends in Stability Testing, with Emphasis on Stability During Distribution and Storage. Drug Development and Industrial Pharmacy 1999, 25 (7), 857–868. https://doi.org/10.1081/DDC-100102246.

14. Bhuyian1 et al,. AN OVERVIEW: STABILITY STUDY OF PHARMACEUTICAL PRODUCTS AND SHELF LIFE PREDICTION 2015 (Volume: 2 Issue: 6). https://www.researchgate.net/publication/285591270_AN_OVERVIEW_STABILITY_STUDY_OF_PHARMACEUTICAL_PRODUCTS_AND_SHELF_LIFE_PREDICTION.

15. Stability Testing of Pharmaceutical Products. J App Pharm Sci 2021. https://doi.org/10.7324/JAPS.2012.2322.

16. Blessy, M.; Patel, R. D.; Prajapati, P. N.; Agrawal, Y. K. Development of Forced Degradation and Stability Indicating Studies of Drugs—A Review. Journal of Pharmaceutical Analysis 2014, 4 (3), 159–165. https://doi.org/10.1016/j.jpha.2013.09.003.

17. Sanjay Bajaj, Dinesh Singla and Neha Sakhuja. Stability Testing of Pharmaceutical Products. Journal of Applied Pharmaceutical Science 02 (03); 2012: 129–138. ISSN: 2231-3354. https://japsonline.com/admin/php/uploads/409_pdf.pdf

18. Jakob, C. A.; Burda, P. Quality Control in Biosynthetic Pathways of N-Linked Glycoproteins in the Yeast Endoplasmic Reticulum. Protoplasma 1999, 207 (1–2), 1–7. https://doi.org/10.1007/BF01294707.

19. Ebrahimi, S. B.; Samanta, D. Engineering Protein-Based Therapeutics through Structural and Chemical Design. Nat Commun 2023, 14 (1), 2411. https://doi.org/10.1038/s41467-023-38039-x.

20. Nag, K.; Islam, Md. J.; Rahman Khan, Md. M.; Rahman Chowdhury, Md. M.; Haq Sarker, Md. E.; Kumar, S.; Khan, H.; Chakraborty, S.; Roy, R.; Roy, R.; Kaunain Oli, Md. S.; Barman, U.; Hasan Bappi, Md. E.; Biswas, B. K.; Mohiuddin, M.; Sultana, N. Development and Qualification of a High-Yield Recombinant Human Erythropoietin Biosimilar; preprint; Bioengineering, 2023. https://doi.org/10.1101/2023.01.22.525046.

21. Nag et al. Satisfying QTPP of Erythropoietin Biosimilar by QbD through DoE-Derived Downstream Process Engineering. Pharmaceutics, MDPI 2023. https://www.mdpi.com/1999-4923/15/8/2087. 10.3390/pharmaceutics15082087.

22. Mamun Al Mahtab et al. Clinical evaluation in adult human revealed the biosimilarity of recombinant Erythropoietin GBPD002 with Eprex^®^. Arch Clin Biomed Res 2023, Fortune Journals. https://www.fortunejournals.com/articles/clinical-evaluation-in-adult-human-revealed-the-biosimilarity-of-recombinant-erythropoietin.pdf. 10.26502/acbr.50170362

23. A Randomized, Double-blinded, Active Controlled Crossover Clinical Trial to Investigate PK, PD and Safety of GBPD002. ClinicalTrials.gov 2022. https://classic.clinicaltrials.gov/ct2/show/NCT05585658

24. ICH Topic Q 5 C: Quality of Biotechnological Products: Stability Testing of Biotechnological/ Biological Products. Annex to the ICH Harmonised Tripartite Guideline for the Stability Testing of New Drug Substances and Products 1996. https://www.ema.europa.eu/en/documents/scientific-guideline/ich-topic-q-5-c-quality-biotechnological-products-stability-testing-biotechnological/biological-products_en.pdf. Accessed on December 20, 2022.

25. Therapeutic Goods Administration. Stability of the finished product. https://www.tga.gov.au/resources/resource/guidance/argom-appendix-2-guidelines-quality-aspects-otc-applications/9-stability-finished-product

26. Chang, Seong-Hun; Kim, Hyun-Jung; Kim, Chan-Wha. Analysis of the Structure and Stability of Erythropoietin by PH and Temperature Changes Using Various LC/MS. Bulletin of the Korean Chemical Society 2013, 34 (9), 2663–2670. https://doi.org/10.5012/BKCS.2013.34.9.2663.

27. Pokharana, M.; Vaishnav, R.; Goyal, A.; Shrivastava, A. STABILITY TESTING GUIDELINES OF PHARMACEUTICAL PRODUCTS. J. Drug Delivery Ther. 2018, 8 (2), 169–175. https://doi.org/10.22270/jddt.v8i2.1564.

28. Wang, W.; Nema, S.; Teagarden, D. Protein Aggregation—Pathways and Influencing Factors. International Journal of Pharmaceutics 2010, 390 (2), 89–99. https://doi.org/10.1016/j.ijpharm.2010.02.025.

29. ICH guidelines. https://www.database.ich.org/sites/default/files/q1a28r22920guideline.pdf. [Last accessed on 2020 Mar 20.

