## Supplementary information for "Determining the shelf life of an erythropoietin alfa biosimilar GBPD002 through stability study"

Table 1: Stability matrix design [Real-time]

| Time | Initial | 3  months | 6  months | 9  months | 12 months | 18 months | 24 months |
| --- | --- | --- | --- | --- | --- | --- | --- |
| Sampling | + + | + + | + + | + – | + – | + – | + – |
| Identification | + + | + + | + + | + – | + – | + – | + – |
| pH | + + | + + | + + | + – | + – | + – | + – |
| Assay | + + | + + | + + | + – | + – | + – | + – |
| *In vitro* assay | + + | + + | + + | + – | + – | + – | + – |
| *In vivo* assay | + + | + + | + + | + – | + – | + – | + – |
| Product-related impurities | + + | + + | + + | + – | + – | + – | + – |
| Process-related impurities | + + | + + | + + | + – | + – | + – | + – |
| Appearance | + + | + + | + + | + – | + – | + – | + – |
| Sterility | + + | + + | + + | + – | + – | + – | + – |
| Endotoxin | + + | + + | + + | + – | + – | + – | + – |

**(++) indicates both real-time and accelerated study to be performed whereas (*+ –*) indicates real-time study to be performed only.*

Table 2: Sampling plan for stability study [Real-time and accelerated]

| Study conditions | Test Frequency | 3  months | 6  months | 9  months | 12  months | 18  months | 24  months |
| --- | --- | --- | --- | --- | --- | --- | --- |
| Real time study samples | | | | | | | |
| 5 ± 3 °C | 1^st^ batch | 4 SBC | 4 SBC | 4 SBC | 4 SBC | 4 SBC | 4 SBC |
|  | 2^nd^ Batch | 4 SBC | 4 SBC | 4 SBC | 4 SBC | 4 SBC | 4 SBC |
|  | 3^rd^ Batch | 4 SBC | 4 SBC | 4 SBC | 4 SBC | 4 SBC | 4 SBC |
| Accelerated study samples | | | | | | | |
| 25 ± 2 °C and  60 ± 5 % RH | 1^st^ batch | 4 SBC | 4 SBC | – | – | – | – |
|  | 2^nd^ Batch | 4 SBC | 4 SBC | – | – | – | – |
|  | 3^rd^ Batch | 4 SBC | 4 SBC | – | – | – | – |

**SBC= Sterile biocompatible container.*

Table 3: EC_50_ value of Erythropoietin alfa concentrated solution

| Batch No. | EC_50_ | | | | | | | |
| --- | --- | --- | --- | --- | --- | --- | --- | --- |
|  | Initial | Accelerated Storage Condition | | Real-time Storage  Condition | | | | |
|  |  | 3M | 6M | 3M | 6M | 9M | 12M | 18M |
| Reference | 0.1789 | 0.1950 | 0.1950 | 0.1605 | 0.1950 | 0.1965 | 0.1904 | 0.1173 |
| Batch 01 | 0.0609 | 0.0868 | 0.1035 | 0.0855 | 0.1035 | 0.0854 | 0.0727 | 0.0737 |
| Batch 02 | 0.1173 | 0.2503 | 0.2126 | 0.1173 | 0.2126 | 0.0680 | 0.0759 | 0.0739 |
| Batch 03 | 0.0687 | 0.0703 | 0.07552 | 0.0687 | 0.0755 | 0.1124 | 0.1346 | 0.0681 |

Table 4: EC_50_ value of GBPD002 4000 IU/mL

| Batch No. | EC_50_ | | | | | | | |
| --- | --- | --- | --- | --- | --- | --- | --- | --- |
|  | Initial | Accelerated Storage Condition | | Real-time Storage Condition | | | | |
|  |  | 3M | 6M | 3M | 6M | 9M | 12M | 18M |
| Reference | 0.2059 | 0.0793 | 0.0920 | 0.1495 | 0.1655 | 0.1640 | 0.2781 | 0.1269 |
| GBFP21013 | 0.0955 | 0.0954 | 0.1280 | 0.1244 | 0.1134 | 0.1098 | 0.1336 | 0.0588 |
| GBFP21014 | 0.2126 | 0.0876 | 0.0889 | 0.0786 | 0.1844 | 0.1639 | 0.1701 | 0.1222 |
| GBFP21015 | 0.0755 | 0.0685 | 0.0685 | 0.0899 | 0.0892 | 0.0922 | 0.1241 | 0.1439 |

Table 5: EC_50_ value of GBPD002 10000 IU/mL

| Batch No. | EC_50_ | | | | | | | |
| --- | --- | --- | --- | --- | --- | --- | --- | --- |
|  | Initial | Accelerated Storage Condition | | Real-time Storage Condition | | | | |
|  |  | 3M | 6M | 3M | 6M | 9M | 12M | 18M |
| Reference | 0.3193 | 0.1676 | 0.1592 | 0.1804 | 0.2103 | 0.1695 | 0.1943 | 0.0923 |
| GBFP21016 | 0.1628 | 0.0984 | 0.0897 | 0.1107 | 0.1078 | 0.1097 | 0.0945 | 0.0868 |
| GBFP21017 | 0.1914 | 0.0761 | 0.1711 | 0.0820 | 0.0806 | 0.0796 | 0.0869 | 0.1439 |
| GBFP21018 | 0.0790 | 0.0941 | 0.1542 | 0.1013 | 0.1121 | 0.1264 | 0.1363 | 0.0844 |

Table 6: EC_50_ value of GBPD002 2000 IU/0.5mL

| Batch No. | EC_50_ | | | | | | | |
| --- | --- | --- | --- | --- | --- | --- | --- | --- |
|  | Initial | Accelerated Storage Condition | | Real-time Storage Condition | | | | |
|  |  | 3M | 6M | 3M | 6M | 9M | 12M | 18M |
| Reference | 0.1605 | 0.2124 | 0.1889 | 0.2083 | 0.1555 | 0.1872 | 0.1232 | 0.1439 |
| GBFP21019 | 0.0855 | 0.0983 | 0.0966 | 0.0749 | 0.0672 | 0.0972 | 0.0976 | 0.0696 |
| GBFP21020 | 0.2126 | 0.1245 | 0.1237 | 0.0596 | 0.0638 | 0.0863 | 0.0790 | 0.0842 |
| GBFP21021 | 0.0931 | 0.0949 | 0.0959 | 0.1087 | 0.1039 | 0.1030 | 0.0595 | 0.0837 |

Table 7: EC_50_ value of GBPD002 5000 IU/0.5mL

| Batch No. | EC_50_ | | | | | | | |
| --- | --- | --- | --- | --- | --- | --- | --- | --- |
|  | Initial | Accelerated Storage Condition | | Real-time Storage Condition | | | | |
|  |  | 3M | 6M | 3M | 6M | 9M | 12M | 18M |
| Reference | 0.1173 | 0.1661 | 0.1375 | 0.1661 | 0.0930 | 0.0774 | 0.0971 | 0.0923 |
| GBFP21022 | 0.0837 | 0.2958 | 0.1797 | 0.1314 | 0.0755 | 0.0752 | 0.0600 | 0.1317 |
| GBFP21023 | 0.0944 | 0.2997 | 0.2007 | 0.1021 | 0.0673 | 0.0830 | 0.0797 | 0.1222 |
| GBFP21024 | 0.0710 | 0.3098 | 0.1740 | 0.0977 | 0.0599 | 0.0632 | 0.0637 | 0.0589 |
